# ALK/ATR combination therapy is effective in neuroblastoma mouse tumors driven by MYCN

**DOI:** 10.1101/2025.07.17.665300

**Authors:** Marcus Borenäs, Dan Lind, Adam Lehnberg, Edit Zenténius, Matilda Esselius, Eva Jennische, Joel Johansson, Jikui Guan, Agata Aniszewska, Jimmy Van den Eynden, Bengt Hallberg, Ruth H. Palmer

**Affiliations:** Department of Medical Biochemistry and Cell Biology, Institute of Biomedicine, Sahlgrenska Academy, University of Gothenburg, Gothenburg, Sweden; Department of Human Structure and Repair, Ghent University, Ghent, Belgium; Cancer Research Institute Ghent (CRIG), Ghent, Belgium

**Keywords:** MYCN, Anaplastic lymphoma kinase (ALK), ATR, Neuroblastoma, lorlatinib, BAY 1895344, elimusertib

## Abstract

One of the key features of high-risk neuroblastoma (NB) is *MYCN*-amplification. While MYCN is still regarded as therapeutically challenging despite intensive efforts to find targeting compounds, inhibitors against Anaplastic Lymphoma Kinase (ALK) are now being evaluated among ALK-driven NB patients with promising results. There is a pressing need to find alternative treatment regimens for *ALK* naïve *MYCN*-amplified patients, as current regimes are accompanied by significant mortality and morbidity. Here we show that genetically removing *Alk* in the *Th-MYCN*-driven engineered NB mouse model (GEMM) resulted in decreased tumor penetrance, and survival of *Th-MYCN* homozygote GEMMs, suggesting that Alk activity contributes to aggressive neuroblastoma development in this model. Given the high levels of replication stress induced in *Th-MYCN* tumors, we employed inhibitors of Ataxia Telangiectasia Rad3 related (ATR), combining ATR and ALK inhibition in a 14-day treatment regime, and comparing with ATR inhibitor monotreatment. *Th-MYCN* tumor bearing mice that received ATRi/ALKi combination treatment exhibited significantly increased survival compared to ATRi alone, that was sustained over time. Together, our data highlight a potentially effective treatment option for the currently challenging category of *MYCN*-amplified NB patients lacking *ALK* mutations.

## Introduction

Approximately 20% of neuroblastoma (NB) patients display amplification of *MYCN*, a hallmark of high-risk (HR) NB, at primary diagnosis ^1–4^. HR-NB has a 5-year overall survival of 50% despite advances in treatment and accounts for around 15% of all childhood cancer deaths ^5, 6^. *MYCN*, together with the *c-MYC* and *MYCL* paralogs, constitute the *MYC* proto-oncogene family which act as transcription factors regulating vital cellular processes such as differentiation, cell growth and programmed cell death ^7–9^. Despite intensive efforts, targeting MYCN in the clinic still represents a challenge, and although clinical trials are ongoing with agents that may impact MYCN (https://clinicaltrials.gov/study/NCT03936465), it is practically regarded as an undruggable target in pediatric NB^10^. New effective treatments for MYCN-driven disease that provide both a durable prolonged therapeutic response with reduced long-term side effects are an acute unmet need.

An immunocompetent genetically engineered mouse model (GEMM) carrying the human *MYCN* gene expressed under the control of the rat tyrosine hydroxylase (Th) promoter has been extensively used to study MYCN in NB ^11^. The *Th-MYCN* GEMM is a well-characterized NB model, and homozygosity for *Th-MYCN* is associated with time-dependent initiation of tumor growth and complete tumor penetrance ^11, 12^. Historically, the *Th-MYCN* NB GEMM model has played an important role in the discovery and development of novel therapeutic combination treatments for pediatric NB patients, a population with limited numbers available for clinical testing.

Deregulated MYC drives cell proliferation, premature S phase progression and error-prone DNA replication, resulting in DNA replication stress (RS) ^7–9, 13, 14^. *MYCN*amplified tumors express increased levels of genes involved in DNA repair and cell cycle checkpoint pathways. These include DNA damage response (DDR) genes such as ataxia telangiectasia and rad3-related (ATR), and checkpoint kinase 1 (CHK1), which have been shown to contribute to tolerance of replication stress ^13–15^. Blocking the DDR may provide significant therapeutic benefit in several malignant diseases such as solid tumors, especially when combining ATR inhibitors with other antitumor agents ^16, 17^. One way to improve outcome in *MYCN*-amplified NB patients would be to exploit the replication stress generated by high levels of *MYCN* expression ^13^. Indeed, previously reported responses to ATR inhibition in mouse NB models support this strategy ^18–23^. Previous work has shown that mis-regulation of the ALK ligand, ALKAL2, in mice potentiates *TH-MYCN* driven NB, without any activating Alk mutations, and that these tumors are sensitive to ALK TKI treatment ^24^. Thus, ALK signaling may impact a wider cohort of NB patients than currently appreciated. In agreement, a recent report identified ALK protein expression in more than 90% of human NB tumors, concluding that all *MYCN*-amplified NBs were ALK-positive, with the majority having high ALK protein expression levels^25^. Preclinical work provides further mechanistic insight, as overexpression of MYCN regulates the transcription of *ALK* ^26^, and ALK activity regulates the initiation of transcription of *MYCN*^27^ and MYCN protein stability ^28–30^.

Here we show that genetic ablation of *Alk* in the Th-MYCN genetically engineered mouse model (GEMM) leads to reduced tumor penetrance and improved survival, implicating ALK activity in MYCN-driven tumor progression. We further investigated the therapeutic impact of combining ATR inhibition with ALK inhibition. A 14-day treatment regimen with ATR and ALK inhibitors resulted in significantly prolonged survival compared to ATR inhibitor monotherapy in Th-MYCN tumor-bearing mice. These findings support a combinatorial approach targeting replication stress and ALK signaling as a potentially effective strategy for MYCN-amplified, ALK–wild-type neuroblastoma.

## Results

### ALK expression in NB

We first confirmed an association between *ALK* and *MYCN* expression in a previously published transcriptomics data set of primary NB (n = 367)^31^. As expected, *ALK* expression was significantly higher in *MYCN* amplified tumors as compared to *MYCN* wild-type tumors (*P* = 2.3e-08; Fig. 1A) and a positive correlation was observed between *MYCN* and *ALK* expression in *MYCN* wild-type (Pearson’s *r* = 0.23; *P* = 5.9e-05: Fig. 1B) but not in *MYCN* amplified tumors. To determine how these findings relate to patient outcome, we performed a survival analysis, comparing patients with the lowest (<P25, i.e., 25^th^ percentile), intermediate (P25-P75) and highest *ALK* (>P75) expression in *MYCN* wild-type and *MYCN* amplified patients. *ALK* expression was negatively correlated to overall survival in *MYCN* wild-type patients (*P* < 0.0051, log rank test; n = 298) and a Cox logistic regression analysis that used the intermediate expression group as a reference population indicated that high *ALK* expression was significantly correlated to worse survival (Hazard Ration (HR) = 1.9; *P* = 0.034; Fig. 1C). Interestingly, this correlation became stronger when more stringent expression thresholds were used (HR = 3.5 above P90 and HR = 4.8 above P95; both *P* < 0.0001; Fig. 1C). In *MYCN* amplified tumors (n = 69) low *ALK* expression was associated with better survival (HR = 0.50; *P* = 0.067 at P25 threshold), an association that became stronger when more stringent thresholds were used (HR dropping from 0.66 at P50 to 0.41 at P05; Fig. 1C). Remarkably, low *ALK* expression had the lowest HR in both *MYCN* amplified and non-amplified patients amongst all receptor tyrosine kinases (RTKs; Suppl. Fig. 1).

**Figure 1.**
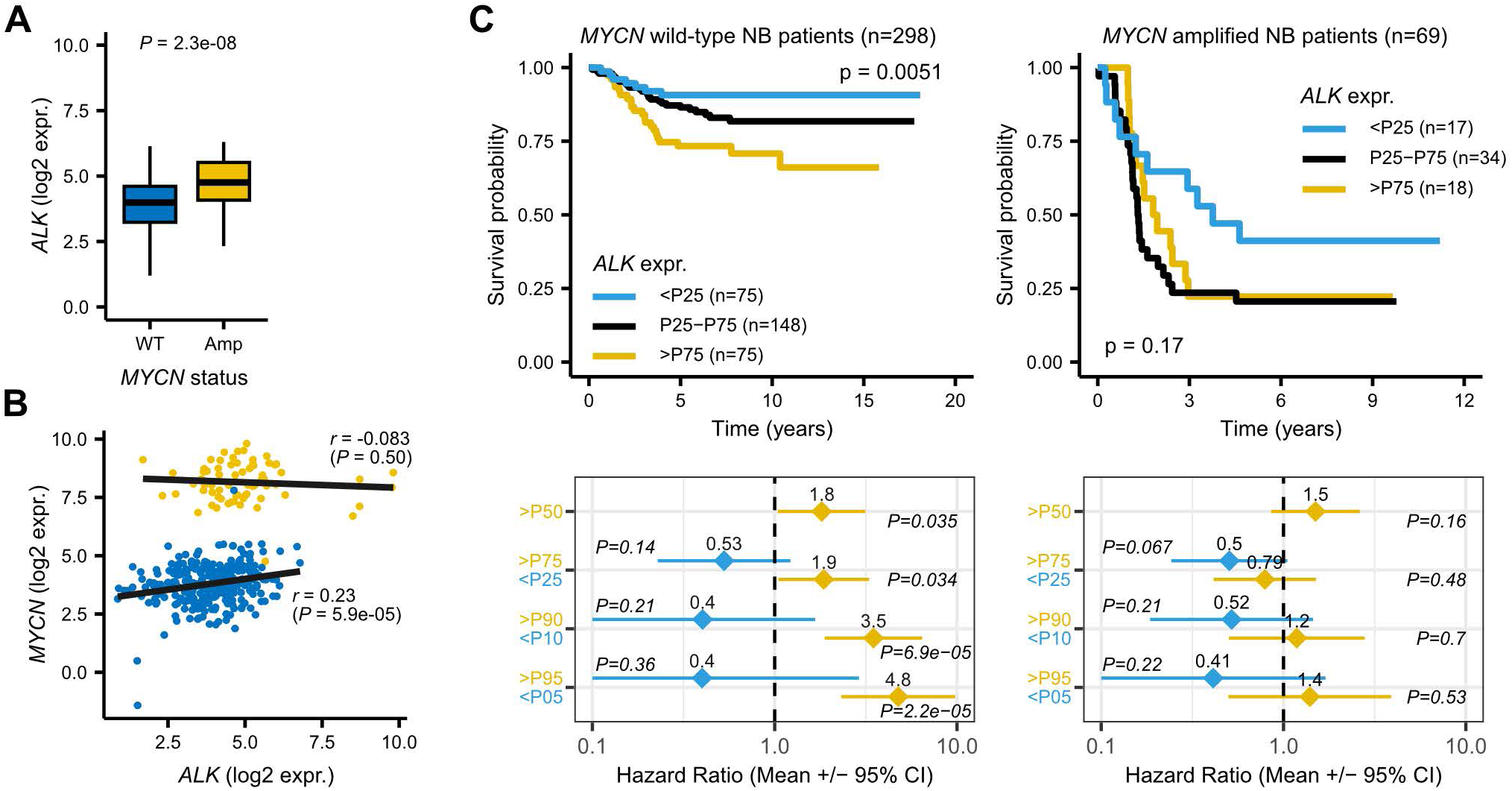
*ALK* dependency in *MYCN* amplified neuroblastoma. **A-C.** Analysis of publicly available primary neuroblastoma (NB) data (n=364)^31^. **A.** Boxplots comparing *ALK* expression (log2 normalized counts) between *MYCN-*amplified (amp) and *MYCN* wild-type (WT) NB tumors. *P* value calculated using two-sided unpaired Wilcoxon’s test. **B.** Correlation plots between *ALK* and *MYCN* expression for *MYCN*-amplified (yellow) and *MYCN* wild-type (blue) tumors. Linear regression line and Pearson’s correlation coefficients indicated. **C.** Upper panels show Kaplan-Meier survival plots comparing overall survival between patients with low (<25^th^ percentile; P25), intermediate (P25-P75) and high *ALK* expression for *MYCN* wild-type and *MYCN*-amplified tumors as indicated. *P* value calculated using log rank test. Bottom panels show corresponding forest plots comparing hazard ratios +/- 95% confidence intervals for different expression thresholds as indicated, as obtained using a Cox proportional hazards regression analysis.

### Genetic removal of *Alk* improves survival of *Th-MYCN*-driven NB in mouse models

These findings indirectly suggest that therapeutically targeting *ALK* could be beneficial in *MYCN* amplified NB patients, even in the absence of *ALK* mutations. To test this clinically highly relevant hypothesis, we focused on *Th-MYCN* driven NB mouse models. We initially explored RNA-seq data from mouse tumors analysed in previous studies^19, 24^, focusing on receptor tyrosine kinase (RTK) expression in RNA-seq data from *Th-MYCN tg/0* and *Alk-F1178S KI/0/Th-MYCN tg/0*. Several RTKs were robustly expressed, including *Ryk*, *Ddr1*, *Aatk*, *Ret*, *Ephb2*, *Epha5/7*, *Lmtk3* and *Alk* (Fig. 2A-B). *Alk* ranked 7^th^ and 5^th^ in the *Th-MYCN tg/0* and *Alk-F1178S KI/0/Th-MYCN tg/0* cohorts respectively. Interestingly, *Alk-F1178S KI/0/Th-MYCN tg/0* tumors exhibited upregulation of the Ret RTK, in keeping with previous reports of transcriptional upregulation of Ret downstream of Ret in NB models ^24, 32^.

**Figure 2.**
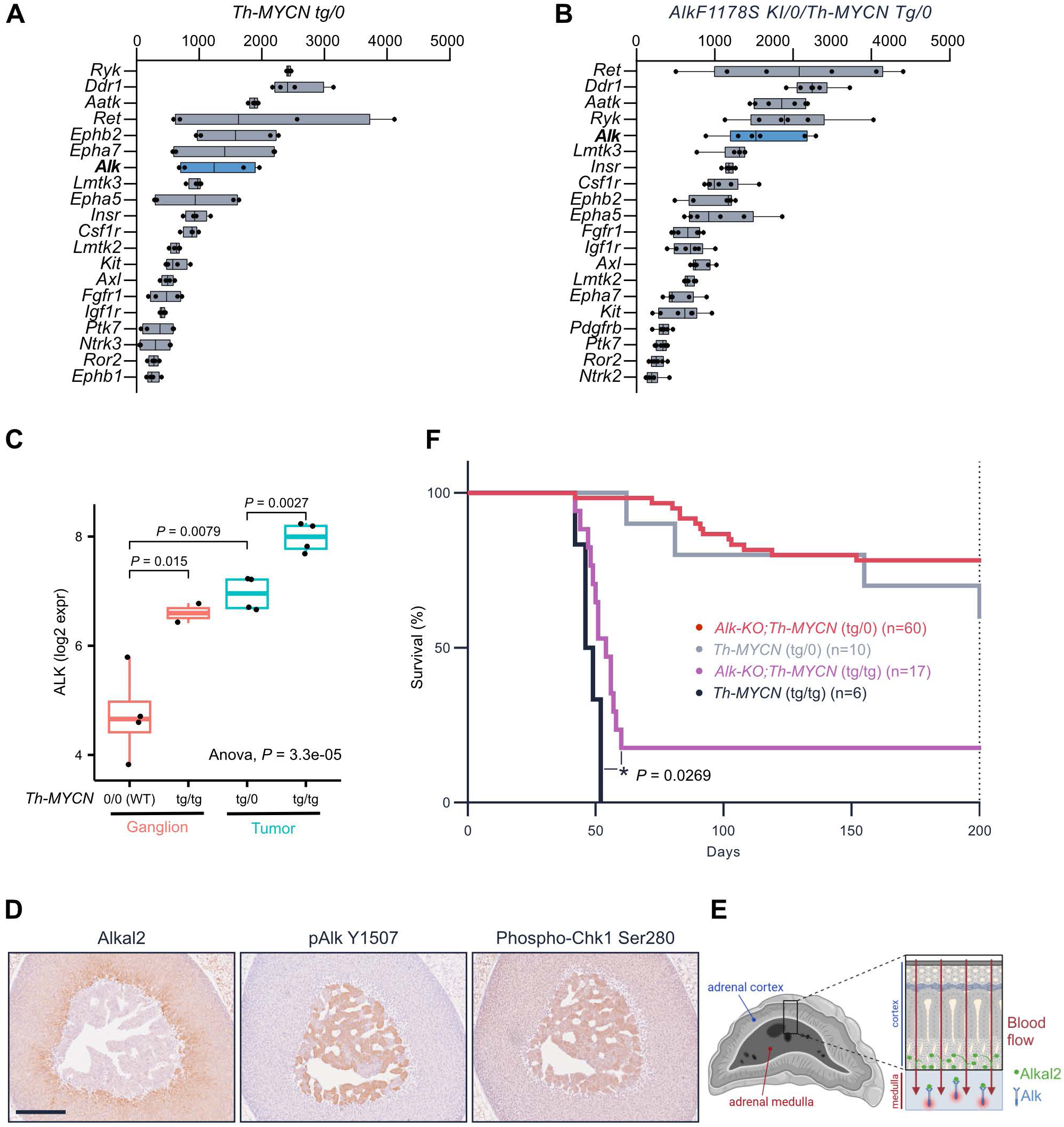
Genetic removal of *Alk* in *MYCN-driven mouse neuroblastoma models*. **A-B.** Boxplots showing expression of the top 20 most abundant RTKs in RNA-seq data from Th-MYCN tg/0 (A) and Alk-F1178S KI/0/ Th-MYCN tg/0 (B) ^19, 24^. **C.** Boxplots comparing *Alk* expression in ganglion and neuroblastoma tissues from *control (wild-type 0/0), Th-MYCN hemizygous (tg/0)* and *Th-MYCN homozygous (tg/tg)* mice, using data from a previous study^33^. Pairwise *P* values calculated using two-sided Student’s t test. **D-E.** Immunohistochemistry of adrenal glands from adult wild-type males (D) and schematic overview (E) of proposed mechanism of Alkal2/Alk/Chk1 signaling in the murine adrenal where Alkal2 is secreted by the adrenal cortex and carried by the blood to the medulla where it activates Alk (created in BioRender). Scale bar indicates 250 µm. **F.** Kaplan-Meier comparing observed survival between different combinations of *Alk (*KO or wild-type) and *Th-MYCN* (hemizygous or homozygous) cohorts, as indicated. Indicated *P* value calculated using log-rank test.

Similar to human tumors, *MYCN* status (i.e., wild-type, hemizygous amplification, homozygous amplification) was associated with *Alk* expression in mouse NB tumors and non-tumoral sympathetic ganglion tissues (*P* = 3.3e-05, ANOVA test; Fig. 2C)^33^. Despite lower expression in control sympathetic ganglion tissues, *Alk* was still clearly expressed (log2 expression values between 3.8 and 5.8, n = 4; Fig. 2C). To confirm Alk expression in normal sympathetic tissues, we performed immunohistochemistry on the adrenal gland, where NB frequently originates in the medulla. Interestingly, and in keeping with single cell RNA-seq data^34, 35^, pY1507-Alk was detected in the adrenal medulla and a complementary expression of the Alk ligand Alkal2 ligand was found in the adrenal cortex (Fig. 2D). This expression pattern suggests that the Alk receptor in the adrenal medulla may be normally activated in a spatially regulated manner by Alkal ligands produced in the cortex and is in keeping with the highly temporospatial regulation of Alk by its ligands in other species (Fig. 2E)^36, 37^.

To determine how these findings correlate with survival, *Th-MYCN* mice were crossed with *Alk* loss-of-function knockouts (*Alk-KO*) ^38^. In agreement with previous reports ^11^, homozygous *Th-MYCN* mice (n=6) rapidly developed aggressive NB and died within two months after birth (Fig. 2F). Remarkably, 18% (3/17) of homozygote *Th-MYCN* mice in an *Alk-KO* background survived for more than 200 days (*P* = 0.0269; log rank test; Fig. 2F). This finding indicates: (1) that Alk may contribute to the development of MYCN-driven NB in the absence of somatic mutations, and (2) that *MYCN*-driven NB may be sensitive to therapeutic intervention with ALK TKIs.

### *Th-MYCN*-driven NB mouse models respond to ALKi/ATRi combination treatment

Previous findings indicate that *MYCN*-driven NB is not just sensitive to ALK inhibition, but also to ATR and CHK1 inhibition ^18–20, 24, 39, 40^. In addition, the ATR downstream target kinase CHK1 has been shown to be upregulated in *MYCN*-amplified NB ^41^, and phosphorylation of Chk1 on S280 is readily detected in adrenal glands from these animals immunohistochemistry (Fig. 2D). We confirmed significantly higher *ATR* expression in *MYCN*-amplified as compared to *MYCN* wild-type human NB tumors (*P* = 1.2e-08; Wilcoxon test; Fig. 3A). Additionally, *MYCN* and *ATR* expression were strongly correlated within the *MYCN* amplified cohort (Pearson’s *r* = 0.50, *P* = 6.7e-04; Fig. 3B) and a survival analysis on these patients indicated worse survival for patients with the highest (i.e., >P75) ATR expression (HR = 2.5, *P* = 0.0053; Fig. 3C). These findings suggest that a combination of ALK inhibition and ATR inhibition could be a valid strategy to treat ALK wild-type, *MYCN*-amplified tumors, similar to that demonstrated previously in *Alk-F1178S;Th-MYCN* GEMM models ^18, 19^. Moreover, robust expression of Chk1, pChk1-S280, pChk1-S317 and pChk1-S345 was observed in both *Alk-F1178S;Th-MYCN* and *Th-MYCN*-driven tumors (Fig. 3D), suggesting high levels of ATR activity. Immunoblotting of *Th-MYCN* and *Alk-F1178S;Th-MYCN* tumors identified pAlk and pChk1 in both tumors, although downstream pAkt and pERK1/2 were stronger in *Alk-F1178S;Th-MYCN* tumors (Fig. 3E). Since a 14-day ATR/ALK inhibitor combination cures NB in a *Alk-F1178S;Th-MYCN* driven model, we investigated the effect of this therapeutic approach in *Th-MYCN*-driven GEMM NB tumors. Tumor-bearing mice were treated with a 14-day regimen that combined the ATR inhibitor elimusertib with the ALK inhibitor lorlatinib; 3 d (days 1-3) elimusertib twice daily, 4 d (days 4-7) lorlatinib twice daily, 3 d (days 8-10) combination, 4 d (days 11-14) lorlatinib and compared with monotherapy with elimusertib; 3 d (days 1-3) elimusertib twice daily, 3 d (days 8-10) elimusertib twice daily, lorlatinib; 14 d twice daily and vehicle; 14 d twice daily ^18, 19^ (Fig. 3F). After treatment, mice were maintained free from therapeutic intervention and tumor volume monitored over time with ultrasound. All mice tolerated the treatment regimen, and no or minimal (<10 mm^3^) tumors were detected after either 14 d combined treatment or monotherapy with elimusertib (Fig. 3F-G). In contrast, vehicle treated homozygote *Th-MYCN* mice displayed rapid progression with only two out of five treated mice reaching the 7 d control (Fig.3F). Interestingly, monotherapy with lorlatinib resulted in a statistically significant (*P=*0.019) improved response to therapy at 7 d with three out of five mice reaching the time point and also continuing to the 14 d control (Fig. 3F). Tumor regrowth was only observed in one of the four homozygous *Th-MYCN* GEMM mice and in two of the five hemizygous *Th-MYCN* that received combination treatment. Five *Th-MYCN* mice that received combination treatment were tumor free for 200 days after treatment cessation (Fig. 2H). In contrast, mice treated with elimusertib as monotherapy (n=4) relapsed rapidly after treatment cessation. These results suggest that an ATR/ALK inhibitor combination may provide therapeutic benefit to a wider group of HR *MYCN*-driven NB patients than previously appreciated. To investigate the potential applicability of ATR/ALK inhibitor treatment among NB patients and whether pCHK1 S280 could be a potential biomarker for ATR/ALK inhibitor sensitivity, we investigated expression of ALK and pCHK1 S280 in commercially available NB samples (Fig 3I-J). A positive correlation between pCHK1 S280 and ALK expression was observed in NB samples, further supporting this site as a downstream marker of ALK activity ^18, 19, 42^.

**Figure 3:**
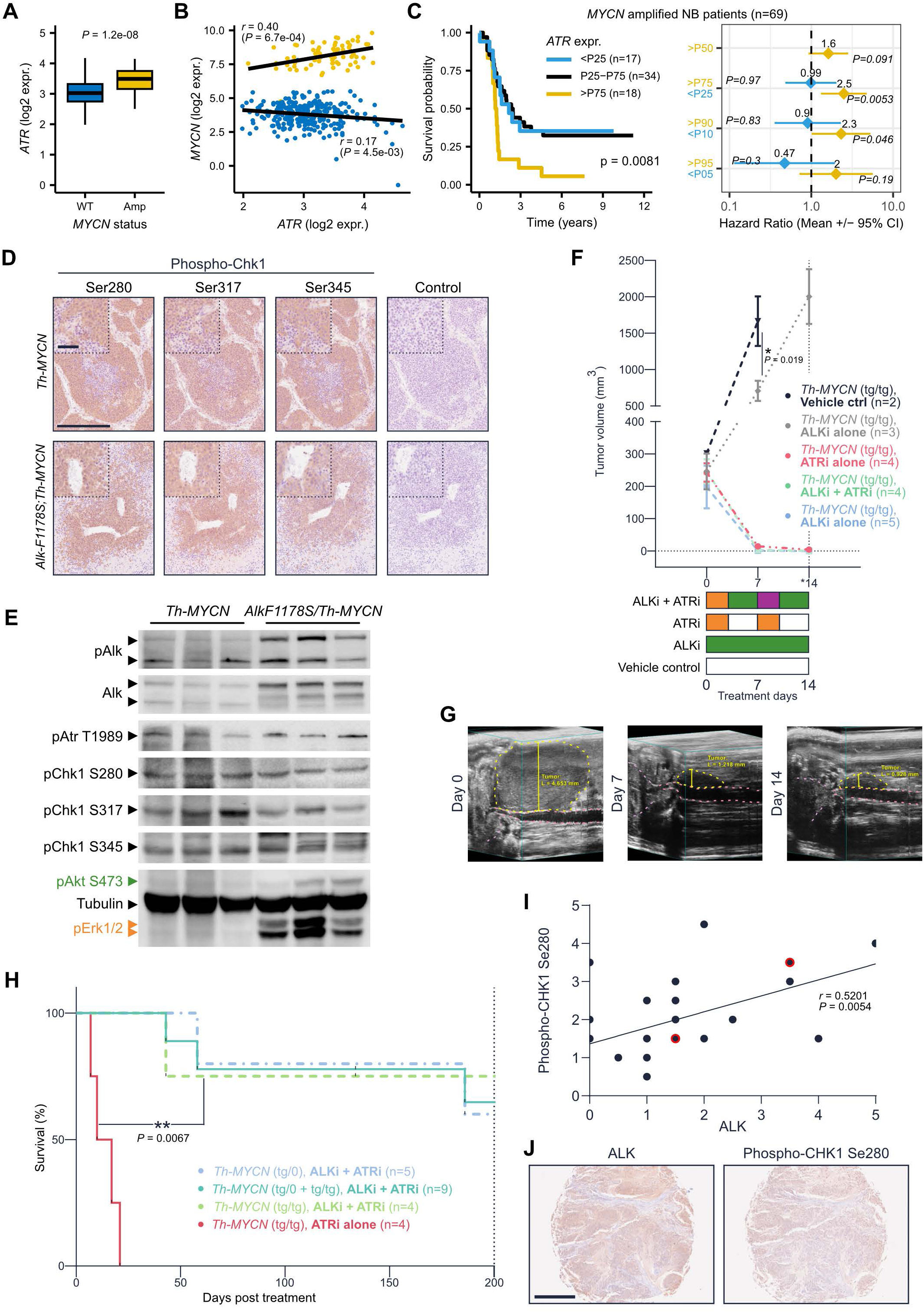
*ALK* wild-type *MYCN* amplified neuroblastoma response to combined ATR/ALK inhibition. **A-C.** Analysis of *MYCN*, *ATR* and survival correlations in publicly available primary NB data (n=364)^31^, similar to Fig. 1A-C. **D.** Representative *Alk-F1178S KI/0/Th-MYCN tg/0* and *Th-MYCN tg/0* tumors were examined for pChk1 S280, S317 and S345. Scale bar indicates 250 µm and 50 µm (in inserts). **E.** Immunoblottning for phosphorylation of Chk1 on sites: S280, S317 and S345, as well as total and pAlk, pAtr, pErk1/2 and pAkt in three *Alk-F1178S KI/0 / Th-MYCN tg/0* and three *Th-MYCN tg/tg* tumors. Tubulin was employed as loading control. **F.** Tumor volume response in *Th-MYCN* mice treated with a 14-day ALKi/ATRi combination (elimusertib + lorlatinib) or monotherapy regimes (elimusertib only, lorlatinib only or vehicle). Combination therapy: 3d elimusertib 25 mg/kg twice/day (orange), followed by 4d lorlatinib 10 mg/kg twice/day (green), followed by 3d elimusertib 25 mg/kg and lorlatinib 10 mg/kg twice/day (purple) and finally 4d lorlatinib 10 mg/kg twice/day (green) after which animals were released from treatment (arrow). Elimusertib 25 mg/kg as monotherapy was given twice per day during the corresponding days of elimusertib in the combination therapy. Lorlatinib 10 mg/kg as monotherapy, or vehicle control, was given twice per day continuously during the 14-day treatment period. Mean ± SD. *=(day 15 and 17 for two of the elimusertib monotreated animals). Unpaired t-test of tumor volume, at d7 between vehicle and lorlatinib treated Th-MYCN (tg/tg) tumor bearing mice, *P=*0.019. **G.** Representativ e ultrasound images of tumor regression during combination treatment in Fig. 2F. Yellow dotted lines denote tumor, pink dotted lines denote aorta and renal artery, kidney outlined with cerise dotted lines. **H.** Kaplan-Meier visualizing survival outcome in treated *Th-MYCN* mice starting (day 0) at treatment release. Teal colored curve, with a ∼65% probability of survival at 200 days, indicates the combined curves of hemizygote *Th-MYCN* (blue dashed line, 60% survival) and homozygote *Th-MYCN* (green dashed line, 75%) mice that received ALKi/ATRi combination treatment. Red colored curve represents *Th-MYCN* homozygote mice that received ATRi monotreatment with elimusertib. Log-rank (Mantel-Cox) test between homozygote *Th-MYCN* (green) that received combination treatment and homozygote *Th-MYCN* (red) that received elimusertib monotreatment, *P=*0.0067. I) Graph displaying simple linear regression for 27 NB samples, duplicate cores, (including 2 ganglioneuroblastoma indicated in red) between ALK and pCHK1 S280 immunohistochemistry stainings. Sections were manually scored (0-5) by an unblinded investigator and the mean of each sample duplicate was used in the tests. Pearson r=0.5201, p=0.0054. J) Example of immunohistochemistry staining for anti-ALK and pCHK1 S280 for one of the duplicate cores among the 27 investigated samples. Scored ALK=5, pCHK1 S280=4. Scale bar indicate 500 µm.

## Discussion

The connection between ALK and MYCN, a known negative prognostic factor when amplified in NB, has been extensively studied, with a strong focus on activating ALK mutations ^43, 44^. The importance of wild type ALK in tumor development is currently of great interest in NB. This study provides compelling evidence for the functional and therapeutic relevance of ALK in MYCN-driven NB, even in the absence of ALK mutations. Consistent with previous reports, we confirmed significantly elevated ALK expression in MYCN-amplified NB tumors. We also show that ALK expression correlates with *MYCN* expression in *MYCN* non-amplified cases and importantly, that elevated ALK expression was associated with worse overall survival in both *MYCN* non-amplified and *MYCN*- amplified patients, with particularly strong prognostic significance in the latter when stringent expression thresholds were applied. These findings are in line with previous observations of ALK protein expression in NB ^25, 45, 46^ and establish ALK expression not only as a biomarker for poor prognosis but also as a potential therapeutic vulnerability in a broader spectrum of NB patients.

It is known that ALK signaling activity regulates MYCN transcription in NB ^27^, and that MYCN directly regulates ALK transcription ^26^, creating a positive feedback loop that potentially enhances tumor growth and survival even in the absence of ALK mutation. We set out to test the hypothesis that wild-type ALK may play a therapeutically significant role in NB, investigating the penetrance of murine MYCN-driven NB in an ALK-deficient genetic background, something that has previously never been tested. Our results show that genetic ablation of *Alk* significantly decreased tumor penetrance in *Th-MYCN* NB bearing mice, demonstrating that ALK contributes to tumorigenesis driven by MYCN, even in the absence of ALK-activating mutations. Our immunohistochemical analysis suggests that the significance of ALK may reflect previously unappreciated spatiotemporal aspects of ALK signaling. Here we show that the ALKAL2 ligand is expressed in the adrenal cortex, while the ALK receptor is expressed and phosphorylated in the adrenal medulla. This striking arrangement likely reflects a physiological function for ALK signaling, where ALKAL2 ligand synthesized in the adrenal cortex, is released into the local capillary and sinusoid network and is ultimately delivered to ALK resulting in ALK signaling activity. In terms of NB development, this may provide a fertile ground for MYCN-driven NB to thrive, coopting physiological ALK signaling processes during normal sympathoadrenal development to enhance tumor survival and growth. As NB can also arise at other sites, it will be important to look at the dynamics of ALK/ALKAL2 expression and activity in the context of neural crest development to address whether co-option of physiological ALK signaling is relevant at other tumor sites, or equally importantly, at metastatic sites. Moreover, the potential for co-option of other RTKs in a similar manner as we show here for ALK will also be important to explore in the future.

Our findings further solidify ATR as an important target in MYCN-amplified NB, which has now been reported by multiple groups ^18–23^. High *ATR* expression correlates with *MYCN* amplification, poorer survival, and we observe robust expression of phosphorylated CHK1 in mouse tumors, suggesting hyperactivation of the ATR/CHK1 DNA damage response axis in this context. Immunoblotting further supported the presence of active ATR signaling in both *Th-MYCN* and *Alk-F1178S;Th-MYCN* tumors. These data provided a rationale for combinatorial inhibition of ALK and ATR in MYCN-driven tumors, which is indeed proves to be effective, even with a short-term 14-day treatment. Perhaps the most striking observation was that sustained tumor remission was significantly improved with ALKi/ATRi combination treatment, with all mice receiving ATR monotherapy relapsing. These findings are particularly significant as they once again extend the therapeutic potential of combination ALKi/ATRi beyond the subset of patients with *ALK* mutations, suggesting broader applicability in high-risk *MYCN-*amplified NB.

The challenge of identifying biomarkers that might identify patients that respond to ALKi/ATRi combination treatment remains. One interesting candidate is CHK1, which has been explored as a therapeutic target in NB^41^. Our data show a positive correlation between *ALK* expression and the ALK signaling dependant pS280 CHK1 site in human NB tumor samples. This further supports the biological interdependence of the ALK and ATR pathways and merits further exploration of pCHK1 as a biomarker that may facilitate the identification of patients most likely to benefit from dual ALK/ATR inhibition.

In conclusion, we show that a 14-d treatment employing the FDA approved ALK inhibitor lorlatinib and the ATR inhibitor elimusertib provide an effective, and gentler, treatment option with less long-term side effects in HR-NB GEMMs. As current treatment options for *MYCN*-driven HR-NB are strenuous and exhausting with significant risk of long-term side effects ^5, 6^, further exploration, perhaps increasing treatment protocols from 14 days to 28 days could be explored as ways to further increase survival rates with the ATRi/ALKi combination. Moreover, these data suggest that pCHK1 may provide a useful biomarker to identify NB cases susceptible to ATR/ALK inhibitor treatment, although further research is needed to confirm these potential findings. Together these results provide attractive therapeutic options for HR *MYCN*-driven NB.

## Supplementary Figure legends

**Supplementary Figure 1. Association between RTK expression and NB patient survival.** A Cox proportional hazards regression analysis was performed comparing NB patients with high (>P75) and low (<P25) gene expression to the intermediate (P25-P75) cohort for *MYCN* wild-type (top panel) and *MYCN* amplified (bottom panel) and for different RTKs as indicated by forest plots. Gene expression for each corresponding RTK indicated by the boxplots on the right of each panel. Only expressed RTKs (i.e., median log2 expression > 1) were considered for the analysis^19, 24^

## Material and methods

### Public data download and analysis

Human NB RNA-seq (batch corrected log normalized counts) and related clinical data (MYCN status, vital state) were obtained from the study of *Cangelosi et al.*^31^ (available at https://www.mdpi.com/2072-6694/12/9/2343/s1). Mice microarray expression data from Murakami-Tonami et al. ^33^ were downloaded from the R2: Genomics Analysis and Visualization Platform (http://r2.amc.nl, R2 identifier: ps_avgpres_gse43419geo20_mm4302).

Statistical analyses were performed with the R statistical package (v4.4). Gene expression correlations were determined using the Pearson correlation method. Gene expression differences between *MYCN* amplified and *MYCN* wild type samples were evaluated using two-sided, two-sample Wilcoxon ranks sum tests. Overall survival analyses of human NB data were performed using the R packages *Survival v.3.7-0* and *Survminer v.0.4.9*. Survival curves were plotted by the Kaplan-Meier method. By default 3 expression groups were compared: high (above P75), intermediate (P25-P75) and low (<P25) gene expression. Log rank tests were performed to assess statistical significance. The Cox proportional hazards regression model was used to calculate hazard ratio’s, always using patients with intermediate gene expression as a reference group. Source code used for public data analyses is available at GitHub https://github.com/CCGGlab/MYCN_ATRi.

### Western blotting and immunohistochemistry

Fresh frozen mouse tumor tissues from three *Alk-F1178S KI/0/Th-MYCN tg/0* (no treatment) and three *Th-MYCN tg/*tg (no treatment) mice were lysed with TissueLyser II (QIAGEN) in RIPA lysis buffer (#89900, ThermoScientific) supplemented with cOmplete™ protease inhibitor cocktail (#11836170001, Roche). Lysates were then clarified by centrifugation and protein concentrations were measured with Pierce™ BCA Protein Assay Kit (#A65453 ThermoScientific). Samples were then boiled in 1x SDS sample buffer, separated with SDS-PAGE and analyzed by immunoblotting. Primary antibodies, including pATR T1989 (#30632), pCHK1 S280 (#2347), pCHK1 S317 (#12302), pCHK1 S345 (#2348), pAKT S473 (#4060), pERK1/2 (#4370) and β-Tubulin (#2146), purchased from Cell Signaling Technology and in-house ALK mAb 135 ^38^ were used to detect protein phosphorylation or total protein levels.

Tumor tissues from one *Alk-F1178S KI/0/Th-MYCN tg/0* (no treatment) and one *Th-MYCN tg/*0 (vehicle treated, 30% PEG 300 (#202371, Sigma-Aldrich) and 2% DMSO (#D4540, Sigma-Aldrich)) were fixed in 10% neutral buffered formalin (HT5011, Merck KGaA, Darmstadt, Germany), embedded in paraffin and sectioned in 5 µm sections using a microtome. After deparaffination, heat mediated antigen retrieval was done with sodium citrate buffer and subsequently treated with 3% H2O2. Normal goat serum 5% was used for blocking. Primary antibodies against CHK1 (#2360, 1:200), pCHK1 S345 (#2348, 1:100), pCHK1 S317 (#12302, 1:1000), pCHK1 S280 (#2347, 1:50) were obtained from Cell Signaling Technology and diluted in in Signalstain antibody diluent (#8112S, Cell Signaling Technology) and incubated in 4°C overnight. PBS was used as primary antibody control. SignalStain Boost IHC Detection Reagent (HRP, Rabbit) (#8114, Cell Signaling Technology) was used for detection, developed in DAB Substrate Kit (#8059, Cell signaling Technology) and counterstained with hematoxylin (#01820, Histolab Products AB). For the Chk1-mouse on mouse staining, SignalStain Boost IHC Detection Reagent (HRP, Mouse) (#8125, Cell Signaling Technology) was used as secondary antibody. Sections were then scanned with Hamamatsu NanoZoomer-SQ Digital slide scanner with 40x magnification.

The adrenal gland disease spectrum tissue arrays (NB642d, Tissue Array, Derwood, MD, USA) were bought from BioCat, Heidelberg, Germany and standard protocol IHC was performed as described above, targeting pCHK1 S280 (#2347, 1:50, Cell Signaling Technology) and ALK (D5F3) (#3633,1:300, Cell Signaling Technology). Duplicate cores for each case.

### Adrenal gland immunohistochemistry

Adrenal glands were fixed in 4 % buffered formaldehyde and embedded in paraffin. Sections (4 µm) were blocked with Blocking reagent (Roche, 11096176001) and incubated overnight with anti-Alkal2 (Inhouse antibody^24^, 1:800), anti-pAlk Y1507 (Abcam, ab73996, 1:400) or anti-pChk1 (Ser 280) (Cell signaling 2347, 1:100). Prior to blocking, heat induced antigen retrieval with Tris/EDTA buffer pH 8.00 (anti-Alkal2) or Citrate buffer pH 6.00 (anti-pAlk and anti-pChk1) was performed. HRP-labeled anti-rabbit antibodies were used as secondary reagent (Jacksson, 711-035-152) and immunoreactions were visualized using Liquid DAB+ substrate chromogen system (Dako K3468). Sections were counter stained with hematoxylin to visualize nuclei, mounted and scanned in a NanoZoomer-SQ (Hamamatsu). Images were subsequently cropped in Affinity designer 2 (2.5.5).

### Crossing of *AlkKO* and *Th-MYCN* and tumor monitoring of offspring

Alk kinase knockout mice (*AlkKO*) mice ^38^ were crossed with *Th-MYCN* ^11^, both on a on a *129X1/SvJ* background, to generate Alk KO/+;Th-MYCN 0/0, Alk KO/KO; Th-MYCN 0/0, Alk KO/+;Th-MYCN tg/0, Alk KO/KO; Th-MYCN tg/0, Alk KO/+; Th-MYCN tg/tg, Alk KO/KO; Th-MYCN tg/tg, Alk +/+; Th-MYCN tg/0 and Alk +/+; Th-MYCN tg/tg. To increase the number of Alk +/+; Th-MYCN tg/0 and Alk +/+; Th-MYCN tg/tg Th-MYCN tg/0 were crossed with Th-MYCN tg/0. Offspring was followed for at least 200 days and end points were defined as when the study ended, the spontaneous death of a mouse or the mouse displaying symptoms that required euthanization of the animal. At end points animals were dissected and tumor presence examined. In total, 198 mice were generated from the *Alk-KO;Th-MYCN* breeding. 13 were excluded due to having been in breeding during the study, 4 litters (19 mice) where most of the animals had not been examined for tumors at sacrifice were excluded, 76 mice were excluded due to lack of the *Th-MYCN* transgene and 2 mice were excluded due to either missing genotype or uncertain tumor status at sacrifice.

In the case of a litter being born over a couple of days, the last day was chosen as the date of birth for the whole litter group. One mouse that developed a tumor had time of sacrifice with an accuracy of a couple of days, therefore the midtime of the period was chosen as the day of sacrifice. 88 mice were included in the study plus an additional five mice, as described below, bringing the total number of mice in the tumor penetrance study to 93.

### ATR and ALK inhibitor treatment and subsequent relapse monitoring

*Th-MYCN tg/0* mice were bred against wild type mice and T*h-MYCN tg/0* to produce Th-*MYCN tg/0* and *Th-MYCN tg/tg* offspring. After weaning, mice were monitored with a Vevo 3100 Imaging System (Fujifilm VisualSonics, Toronto, Canada) ultrasound and once a tumor was detected and reached a volume of between 150 and 320 mm^3^ treatment was initiated. Treatment consisted of a tailormade lorlatinib (RLor-10-21, Reagency, Melbourne, Australia) (10 mg/kg twice daily when given) and elimusertib (HY-101566, MedChemTronica, Sollentuna, Sweden) (25 mg/kg twice daily when given) combination treatment regime for a total of 14 days or monotherapy with elimusertib ^18, 19^. The inhibitors were diluted in 30% PEG 300 (#202371, Sigma-Aldrich) and 2% DMSO (#D4540, Sigma-Aldrich). 3D image acquisition was performed on day 0, day 7 and day 14 (day 15 and 17 for two of the monotreated animals). After the 14-day combination treatment animals were released from treatment and monitored for tumor relapse as defined as growing mass reaching at least 100 mm^3^. Image acquisition, volume calculations, treatment regime, ultrasound screening and tumor monitoring were performed in the same manner as previously described ^18^. Five Th-MYCN tg/tg mice that received treatment (3 received combination treatment and 2 received control treatment) were also included in the *Alk-KO;Th-MYCN* penetrance study. 10 days were added to the date of treatment initiation to estimate, by empirical experience, when the mouse might have displayed symptoms of tumor burden had treatment not been given.

### Mouse husbandry and genotyping

Genotyping for Th-MYCN was performed with following primers: 5′-AGGGATCCTTTCCGCCCCGTTCGTTTTAA-3′, 5′-TGGAAAGCTTCTTATTGGTAGAAACAA-3′, ACTAATTCTCCTCTCTCTGCCAGTATTTGC-3′ and 5′- 5′- TGCCTTATCCAAAATATAAATGCCCAGCAG-3′ while genotyping for Alk KO was performed utilizing: 5′-CCTGATGATCAGAGCTTC-3′, 5′-CCTGTGGCTCACTTAACT-3′, 5′-ATAGTTCGCCAGCGCGCACCGTA-3′ and 5′-TGCAGTCACTGCGAGGTAGACG-3′. Mice were genotyped by Transnetyx (Cordova, TN). Protocols and procedures performed cohered with the Regional Animal Ethics Committee approval, Jordbruksverket (Id.nr. 003225, Dnr 5.8.18-11644/2020, Id.nr. 004793, Dnr 5.8.18-02319/2023).

## Acknowledgements

This work has been supported by grants from the Swedish Cancer Society (BH: CAN21/1525; RHP: CAN21/1459), the Swedish Research Council (RHP: 2023-02433; BH: 2021-01192), The Wallenberg Foundation (RHP: KAW 2015.0144), the Swedish Childhood Cancer foundation (BH: PR2021-27: RHP: PR2022-0029), Assar Gabrielsson’s Foundation (MB: FB23-71), the Gothenburg Society of Medicine (MB: GLS-986836) and the Research Foundation Flanders (JVdE, RHP: FWO.OPR.2023.0063).

## Contributions

BH and RHP conceived the research project. Wet lab experiments were conducted by MB and DEL with the help from AL, EZ, ME, JJ, JG, and AA. MB, JJ and JVdE conducted the bioinformatics analysis. MB, RHP and BH coordinated the manuscript that was developed with all authors.

## Notes

### Competing Interest Statement

The authors have declared no competing interest.

