## Supplementary figures and images for "ALK/ATR combination therapy is effective in neuroblastoma mouse tumors driven by MYCN"

### Supplemental Fig 1

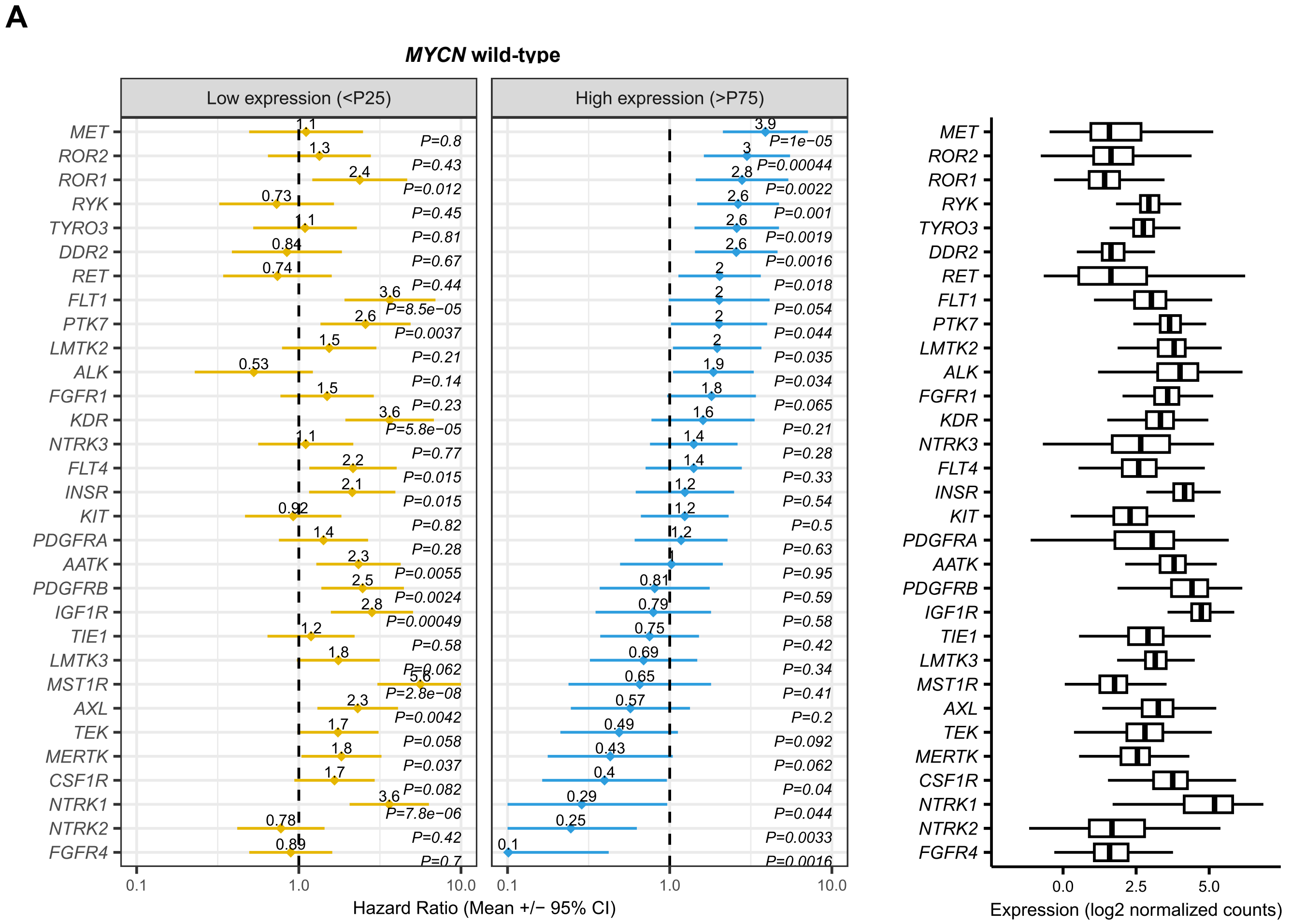

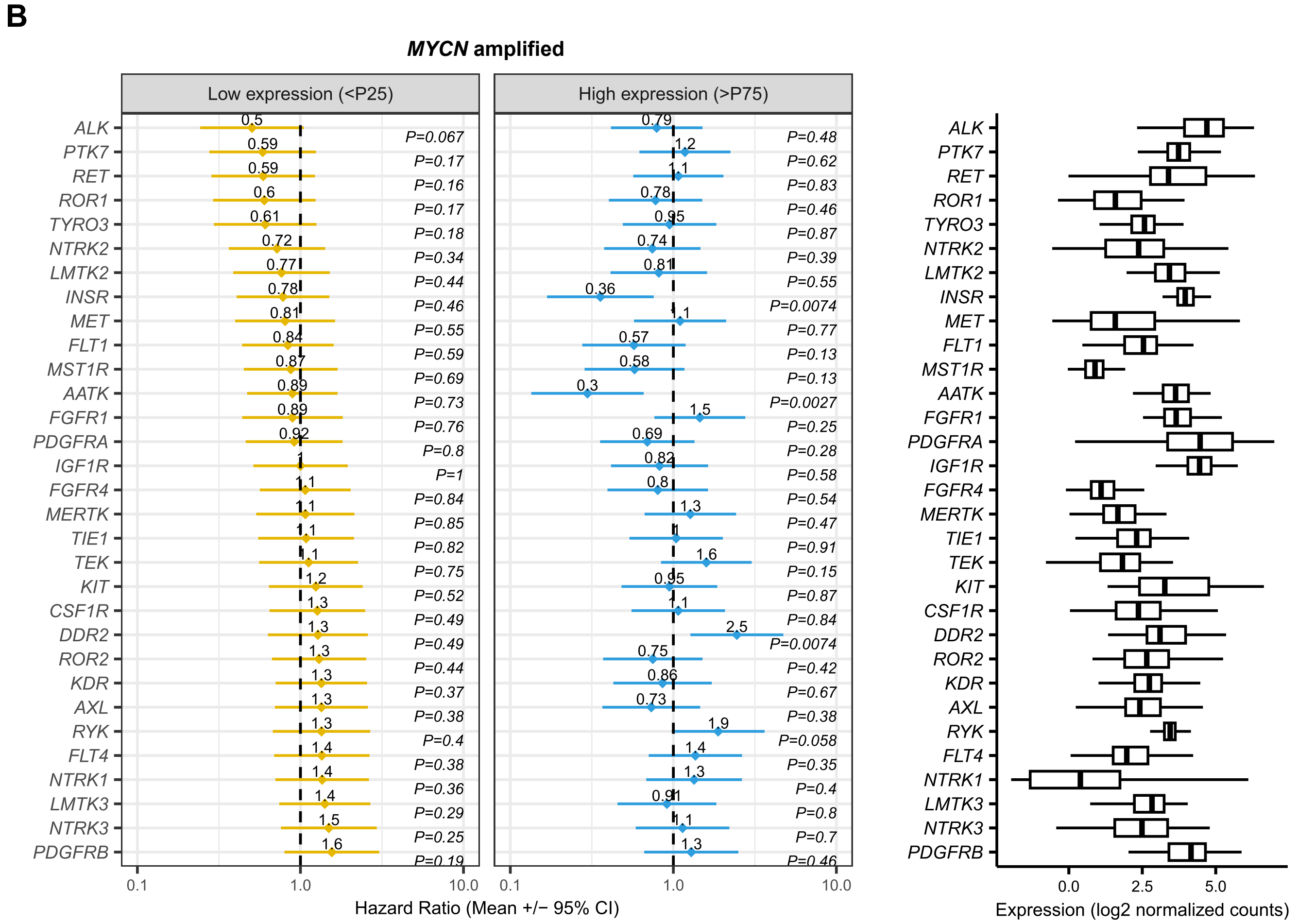
